# Arabidopsis MEB3 functions as a vacuolar transporter to regulate iron accumulation in roots

**DOI:** 10.1101/2023.03.09.531946

**Authors:** Kaichiro Endo, Arpan Kumar Basak, Alwine Wilkens, Mohamadreza Mirzaei, Stanislav Kopriva, Kenji Yamada

## Abstract

Iron is an essential nutrient for plant photosynthesis and development, but excess iron leads to stress. After absorption from the soil, plants store iron in roots and distribute it to shoots via long-distance transport. Vacuole serves as the iron storage organ in root cells, maintaining cellular iron homeostasis, and vacuolar iron transporter (VIT) family proteins have been identified as plant vacuolar iron transporters. However, the contribution of vacuolar iron transporters to the overall iron homeostasis of plants is not fully understood. Here, we show that MEMBRANE PROTEIN OF ER BODY 3 (MEB3), a VIT family member, is a vacuolar iron transporter involved in root–shoot iron distribution in *Arabidopsis thaliana*. Heterologous expression of Arabidopsis *MEB3* in yeast restored the iron resistance phenotype of the vacuolar iron transporter deficient mutant *ccc1*, indicating that MEB3 regulates iron transport. In Arabidopsis, *MEB3* was expressed in almost all tissues, albeit to higher levels in roots and seedlings, and the MEB3 protein localized to the tonoplast. At low iron concentration, *meb3* knockout mutants accumulated less iron in shoots, suggesting that MEB3 promotes iron accumulation in shoots. Consistently, *meb3* mutants exhibited reduced growth compared with the wild type upon transfer to iron-deficient medium. However, at high iron concentration, *meb3* mutants accumulated more iron in shoots and less iron in roots than the wild type, indicating the impairment of proper iron distribution in *meb3* mutants. These findings demonstrate that MEB3 is a vacuolar iron transporter involved in root-to-shoot iron distribution.

## INTRODUCTION

Iron is a heavy metal essential for many processes related to plant growth (Chaffai and Koyama 2011; Andresen et al. 2018), such as photosynthesis (Jeong and Guerinot 2009; Kroh and Pilon 2020), chlorophyll biosynthesis (Mochizuki et al. 2010), metabolic gene expression (López-Millán et al. 2013), and suppression of glycolysis and phloem glucose loading (Thimm et al. 2001). Therefore, the regulation of iron content and distribution in tissues and organs is deeply connected to the survival strategy of plants.

Ferric ions (Fe^3+^) present in soil are reduced to ferrous ions (Fe^2+^) by ferric-chelate reductase found on the surface of root epidermal cells (Robinson et al. 1999; Connolly et al. 2003), and then transported into root cells via the plasma membrane-localized IRON REGULATED TRANSPORTER 1 (IRT1) protein (Vert et al. 2002; Connolly et al. 2002). The incorporated cytosolic iron is then stored in the vacuole of root cells or transported to shoots via the vascular tissue. To enable the long-distance transport of iron from roots to shoots, the metal ions are converted into metal-chelate complexes such as Fe^3+^-citrate, Fe^3+^-mugineic acid, and Fe^2+^-nicotianamine (Fe^2+^-NA) (Kobayashi and Nishizawa, 2012). Iron uptake and translocation to shoots are enhanced by iron deficiency (Schwarz and Bauer, 2020). The expression of *IRT1* and chelator biosynthesis genes in roots is upregulated by iron deficiency in shoots, and it enhances the ability of iron delivery to shoots (Kobayashi et al. 2019). In Arabidopsis (*Arabidopsis thaliana*), IRON MAN/FE UPTAKE-INDUCING PEPTIDES (IMA/FEPs) were recently identified as signaling molecules derived from shoots that positively regulate the expression of *IRT1* and chelator biosynthesis genes in roots, indicating that iron uptake by roots is controlled via long-distance signaling from shoots under iron-deficient conditions (Grillet et al. 2018; Hirayama et al. 2018; Tabata et al. 2022).

In plant cells, iron is distributed to plastids and mitochondria for various applications (Andresen et al. 2018), but excess iron is stored mainly in vacuoles (Lanquar et al. 2010; Donner et al. 2012). Arabidopsis VACUOLAR IRON TRANSPORTER 1 (VIT1), a homolog of yeast (*Saccharomyces cerevisiae*) CCC1, is responsible for vacuolar iron storage in the provascular and endodermal cells of the embryo, which is important for early seed germination (Donner et al. 2012; Grillet et al. 2014; Kim et al. 2006). VIT1 and its homologs possess a conserved multispanning transmembrane region, named DOMAIN OF UNKNOWN FUNCTION 125 (DUF125), and these proteins together form a large VIT family. The VIT family includes Arabidopsis VACUOLAR IRON TRANSPORTER LIKE 1 (VTL1), VTL2, and VTL5, *Lotus japonicus* SEN1, soybean (*Glycine max*) nodulin-21, and Arabidopsis MEMBRANE OF ER BODY 1 (MEB1) and MEB2 (Yamada et al. 2013). Arabidopsis VTL1, VTL2, VTL5, MEB1, and MEB2 proteins exhibit iron transport activity when the corresponding genes are heterologously expressed in yeast (Gollhofer et al. 2014; Yamada et al. 2013). Arabidopsis *VTL1* encodes a vacuolar protein and is expressed in the roots, hypocotyls, and cotyledons of seedlings, suggesting that VTL1 is a major vacuolar iron transporter that regulates iron accumulation in seedlings (Gollhofer et al. 2011). In mature Arabidopsis plants, the expression of *VTL1* is downregulated by iron deficiency, suggesting that plants reduce the level of vacuolar iron transporters to increase the mobilization of vacuolar iron (Yan et al. 2016). The iron transport activity of *L. japonics* SEN1 and soybean nodulin-21 is obscure, but the corresponding genes are expressed in root nodules and are suggested to be involved in nitrogen fixation (Delauney et al. 1990; Hakoyama et al. 2012). Alfalfa (*Medicago truncatula*) and soybean SEN1 homologs (MtVTL8 and GmVTL1, respectively) are believed to function as iron transporters in root nodules (Liu et al. 2020; Brear et al. 2020; Walton et al. 2020).

Besides the VIT family proteins, FERROPORTIN 2 (FPN2) has also been identified as a vacuolar iron transporter (Morrissey et al. 2009), while NATURAL RESISTANCE-ASSOCIATED MACROPHAGE PROTEIN 3 (NRAMP3) and NRAMP4 have been identified as vacuolar iron exporters in Arabidopsis (Lanquar et al. 2005). The expression of *NRAMP3* and *NAMP4* is upregulated in Arabidopsis roots under iron-deficient conditions, indicating that *NRAMP3* and *NAMP4* are responsible for iron mobilization from vacuoles during iron deficiency. Together with the VIT family proteins, these iron importers and exporters contribute to iron mobilization in plants.

Unlike other VIT family proteins, which localize to vacuoles, Arabidopsis MEB1 and MEB2 proteins localize to endoplasmic reticulum (ER) bodies (Yamada et al. 2013). ER bodies are ER-derived, rod-shaped organelles reported in the Brassicaceae family and the closely related Cleomaceae and Capparaceae families (Behnke and Eschlbeck, 1978). ER bodies accumulate β-glucosidases, which hydrolyze glucosinolates, releasing metabolites that repel herbivores (Nakazaki et al. 2019; Yamada et al. 2020) and facilitate root microbiota establishment in Arabidopsis (Hara-Nishimura and Matsushima 2003; Yamada et al. 2009; Basak et al. 2022). The function of MEB1 and MEB2 in glucosinolate metabolism remains obscure, but both proteins exhibit iron and manganese transport activity and play a role in maintaining the rod-shaped morphology of ER bodies (Yamada et al. 2013; Basak et al. 2021).

In this study, we characterized the closest homolog of MEB1 and MEB2 in Arabidopsis, namely, MEMBRANE PROTEIN OF ER BODY 3 (MEB3; AT4G27870). The function of MEB3 is thought to be unrelated to ER bodies because we previously showed that *MEB3* gene expression is not controlled by NAI1, a transcription factor regulating most ER body-related genes in seedlings (Yamada et al. 2013). Here, we investigated the subcellular localization and iron transport activity of MEB3 in Arabidopsis. Our results showed that MEB3 serves as a vacuolar, but not ER body-specific, iron transporter. The *MEB3* gene was expressed mainly in roots, and iron level and plant growth were reduced in Arabidopsis *meb3* knockout mutants under iron-deficient conditions. Our findings suggest that MEB3 plays a crucial role in iron accumulation and mobilization in Arabidopsis root cells and is involved in root-to-shoot iron translocation in response to iron availability.

## RESULTS

### VIT family proteins consists three subfamilies, and MEB3 belongs to the MEB subfamily

Our previous analysis indicated that Arabidopsis possesses three highly homologous proteins, namely, MEB1, MEB2, and At4g27870 protein (Yamada et al. 2013). These proteins have a multispanning transmembrane region that shows homology to VIT family proteins and is proposed to have a metal transporter function. This transmembrane region of VIT family proteins is dubbed DUF125. To estimate the functional similarity of VIT family proteins, we generated an unrooted phylogenetic tree of amino acid sequences harboring DUF125 (Figure 1). The results showed that VIT family proteins could be divided into three subfamilies: VIT1, SEN1/NODULIN21, and MEB. The MEB subfamily contained proteins belonging not only to the Brassicaceae family (e.g., Arabidopsis and *Brassica napus*) but also to other plant families (Figure 1). Arabidopsis MEB1, MEB2, and MEB3 clustered in the MEB subfamily.

**Figure 1.**
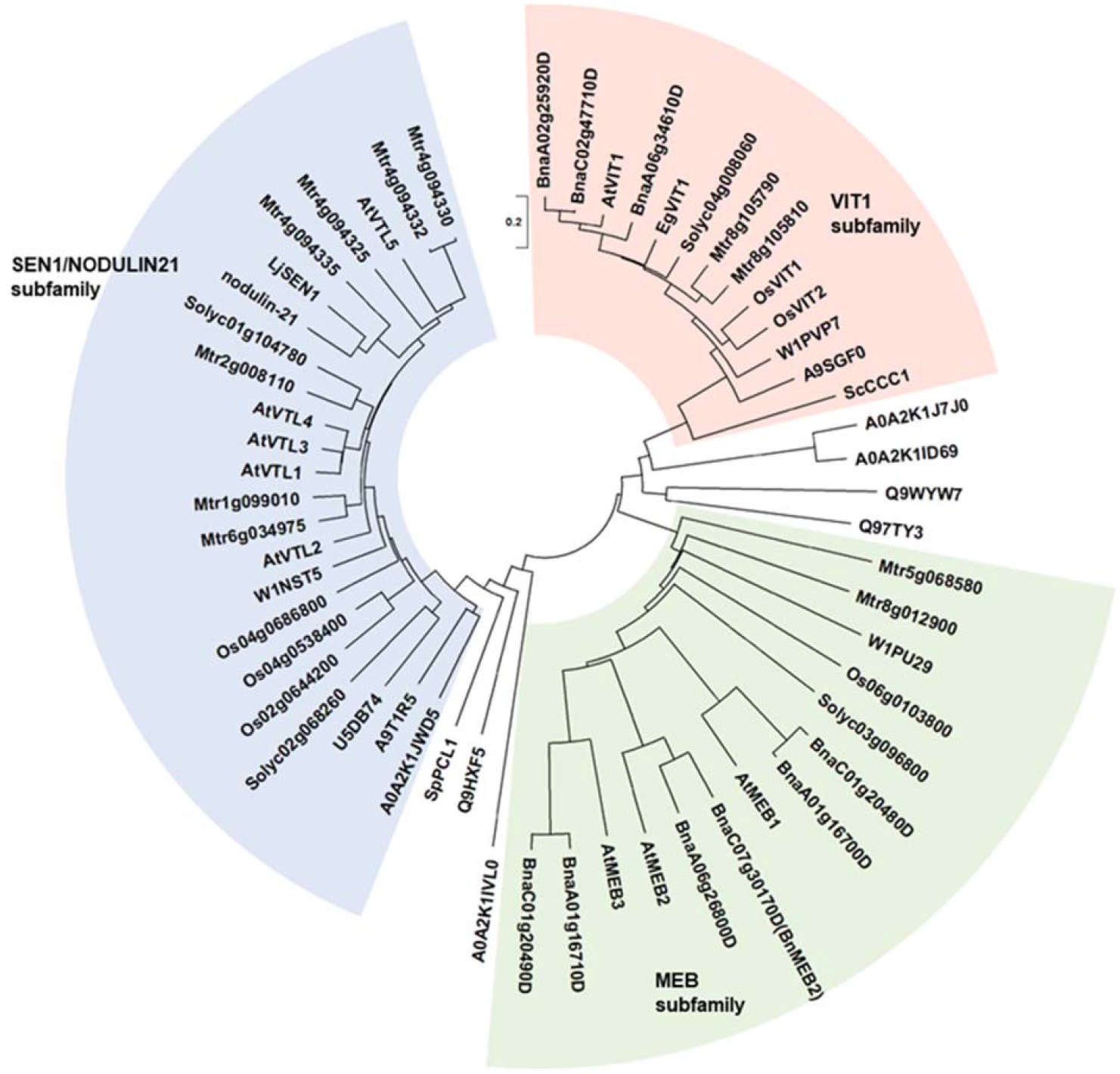
Phylogenetic tree of VIT family proteins. The three subfamilies, VIT1 (red background), SEN1/NODULIN21 (blue background), and MEB (green background), are shown. The proteins used for the phylogenetic analysis are listed in Supplementary Table S1. The scale bar represents the evolutionary distance, expressed as the number of substitutions per amino acid.

The EgVIT1 protein of eucalyptus (*Eucalyptus grandis*) contains four amino acid residues in DUF125, namely, Asp43, Glu72, Met80, and Tyr175, which together form a cavity for transporting iron (Kato et al. 2019). Additionally, amino acid substitution of Gly76 severely affected the iron transporter function of Arabidopsis VIT1 (Mary et al. 2015). Therefore, we investigated the presence of these amino acid residues in VIT family proteins (Supplementary Figure 1). We found that Asp43, Glu72, Met80, Tyr175, and Gly76 residues were well conserved in the VIT1 subfamily, but that Glu72 and Tyr175 were not conserved in the SEN1/NODULIN21 subfamily, and Asp43 and Glu72 were not conserved in the MEB subfamily. Collectively, these results suggest that, in addition to functioning as iron transporters, the VIT family proteins of each subfamily also perform unique functions.

### MEB3 functions as an iron transporter in yeast

To examine whether MEB1–3 proteins exhibit iron transporter activity, we employed a yeast expression system. First, we checked the subcellular localization of MEB proteins in yeast by expressing the *MEB* genes as fusions with the *green fluorescent protein* (*GFP*) gene. The GFP-MEB3 fusion protein showed vacuolar localization, but the vacuolar localization of GFP-MEB1 and GFP-MEB2 fusion proteins was obscure; GFP-MEB1 appeared to form aggregates, and GFP-MEB2 localized to the perinuclear ER membrane in yeast cells (Figure 2A). Next, we used the vacuolar iron transporter deficient yeast mutant *ccc1* and tested whether the *MEB1*–*3* genes could recover the iron-sensitive phenotype of the mutant (Figure 2B). The yeast *ccc1* mutant showed normal growth on synthetic galactose (SGal) medium but reduced growth on SGal containing 3 mM iron, indicating that the *ccc1* mutant was unable to sequester the cytosolic iron in the vacuole, which resulted in the increase of cytosolic iron to a toxic level. Overexpression of *MEB1* and *MEB2* slightly improved the growth of the *ccc1* mutant on iron-containing medium (Figure 2B) (Yamada et al. 2013). The growth of the *ccc1* mutant was restored to a significantly higher level by *MEB3* overexpression than by *MEB1* or *MEB2* overexpression (Figure 2B). Considering the vacuolar localization of MEB3 in yeasts (Figure 2A), these results suggest that MEB3 transports iron into the vacuole to prevent cytosolic iron toxicity in the *ccc1* mutant. Finally, we measured the iron contents of yeast cells cultured in liquid medium containing 0.5 mM iron that allows *ccc1* mutant growth (Figure 2C). We found that overexpression of both *MEB3* and *VIT1* led to iron accumulation in *ccc1* mutant yeast cells, supporting the idea that MEB3 works as a vacuolar iron transporter in the heterologous yeast expression system.

**Figure 2.**
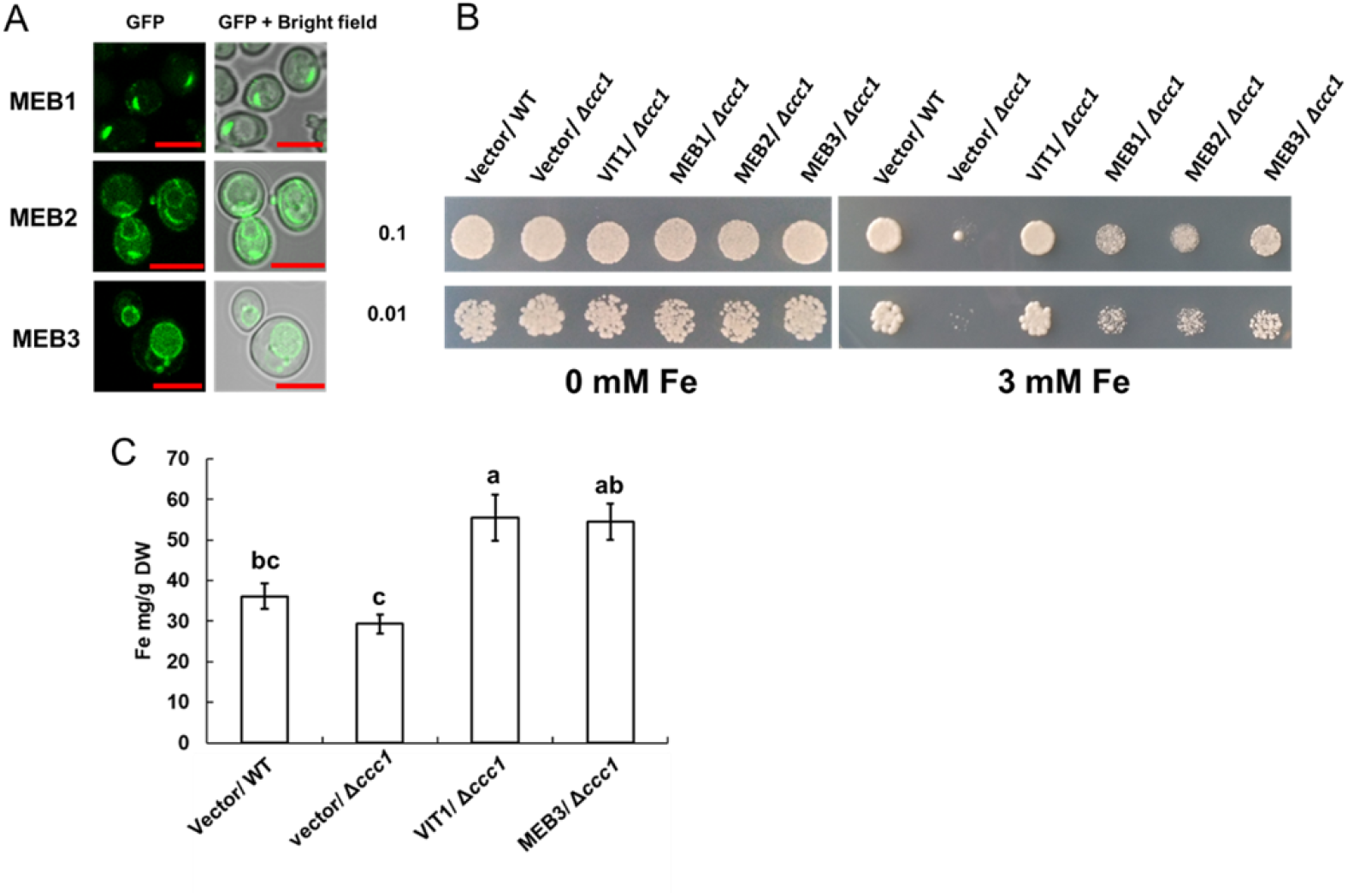
Protein subcellular localization, functional complementation, and iron uptake assays performed using yeast cells. A, Subcellular localization assays of GFP-MEB1, GFP-MEB2, and GFP-MEB3 fusion proteins. Panels show confocal microscopy images (left) and bright field images (right). Scale bars = 10 µm. B, Functional complementation assay. The wild-type (WT) and iron-sensitive Δ*ccc1* strains of yeast were transformed with the empty vector (Vector) or vectors harboring *VIT1*, *MEB1*, *MEB2*, or *MEB3* and grown on SGal medium supplemented with or without 3 mM iron for 4 days. Numbers on the left denote the concentration (OD_600)_ of the spotted yeast cells. C, Iron uptake assay. The chart shows the total iron contents of yeast cells, as determined by the iron colorimetric assay, after 1 day of incubation on medium containing 0.5 mM (NH₄)₂Fe(SO₄)₂. Data represent mean ± standard error (SE; *n* = 3). Different lowercase letters indicate significant differences (*p* < 0.05; Tukey’s test).

### MEB3 is a vacuolar membrane protein

To investigate the subcellular localization of MEB3 *in planta*, we constitutively expressed a fusion protein comprising tandem dimer tomato (tdTOM), a fluorescent protein, and MEB3 (tdTOM-MEB3) in Arabidopsis. Red fluorescence was observed on membrane-bound structures in cotyledons and roots. MEB1 and MEB2 localized to ER body membranes in Arabidopsis (Yamada et al. 2013). However, in plants expressing the *tdTOM-MEB3* fusion, no ER body-like structures could be detected in cotyledons and roots, which are otherwise rich in ER bodies. This indicates that the subcellular localization pattern of MEB3 is different from that of MEB1 and MEB2. Notably, plants expressing *tdTOM-MEB3* showed fluorescence along transvacuolar strands and spherical structures in leaves and roots, respectively, suggesting that tdTOM-MEB3 localizes to the tonoplast (Figure 3A); however, we could not exclude the plasma membrane-localization of MEB3 in this analysis. To investigate the subcellular localization of MEB3 in further detail, we transiently expressed the *GFP-MEB3* fusion in protoplasts isolated from Arabidopsis mesophyll cells (Figure 3B) and tobacco (*Nicotiana tabacum*) BY-2 culture cells (Figure 3C). In both cell types, the GFP-MEB3 fusion protein was detected on the tonoplast but not on the plasma membrane. These data indicate that Arabidopsis MEB3 is a tonoplast protein.

**Figure 3.**
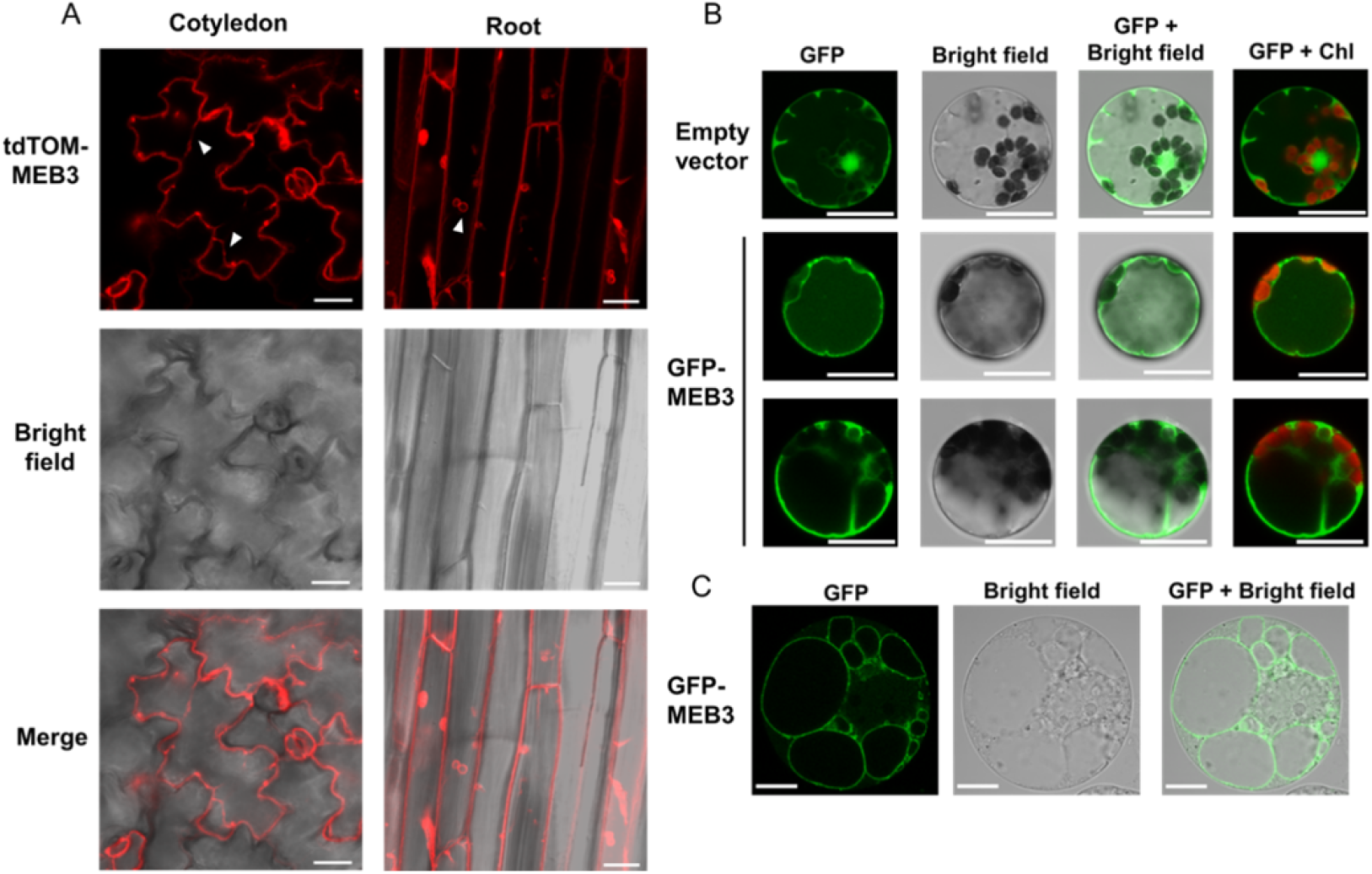
Determination of the tonoplast localization of MEB3 by confocal microscopy. A, The image shows tdTOM fluorescence signal in the epidermal cells of cotyledons and roots of 5-day-old and 14-day-old transgenic plants, respectively. B and C, Cytosolic localization of GFP (Empty vector) or GFP-MEB3 in transiently transformed Arabidopsis (B) and BY-2 (C) protoplasts. ‘Chl’ indicates chlorophyll autofluorescence. Scale bars = 20 µm.

### *MEB3* exhibits tissue-specific expression and mild response to jasmonic acid (JA)

We examined the expression patterns of *MEB1–3* and *VIT1* genes in different plant organs (Figure 4A). All *MEB* genes were highly expressed in the cotyledons and roots of 5- and 14-day-old plants, respectively. The expression level of *MEB3* was higher than that of *MEB1* and *MEB2* in green siliques and flowers. Unlike *MEB3*, the *VIT1* gene showed high expression level in green siliques; however, its expression could not be detected in other organs examined, as reported previously (Kim et al. 2006). These results suggest that MEB3 contributes to iron accumulation in cotyledons, roots, siliques, and flowers.

**Figure 4.**
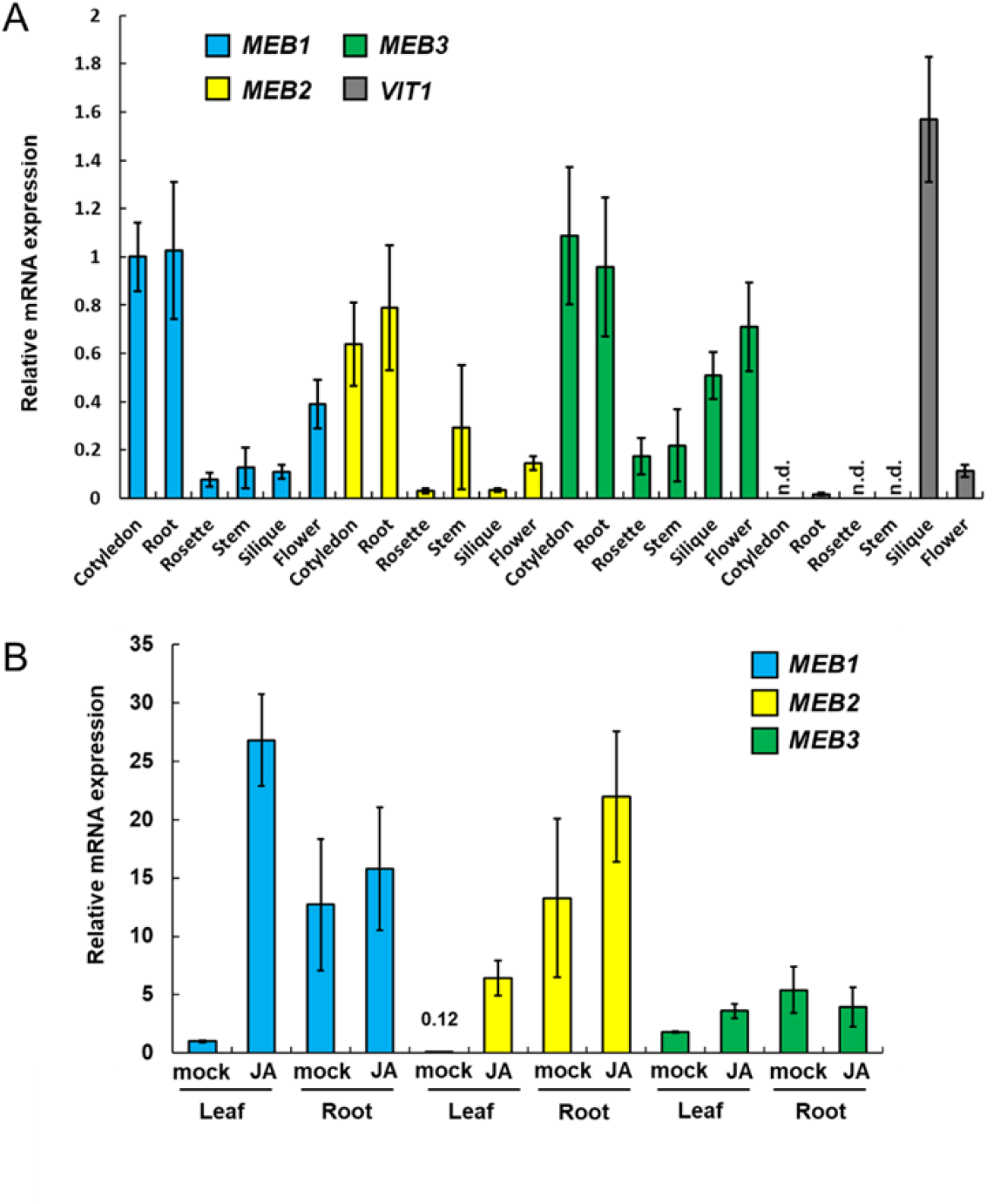
Expression analysis of *MEB1–3* and *VIT1* genes in Arabidopsis. A, Expression of *MEB1*, *MEB2*, *MEB3*, and *VIT1* genes in different organs of Arabidopsis plants. The chart shows gene expressions in the cotyledons of 5-day-old seedlings; rosette leaves and roots of 14-day-old seedlings; and stems, siliques, cauline leaves, and flowers of 6-week-old plants. The transcript level of each gene was normalized relative to that of *UBQ10*. Data represent mean ± SE (*n* = 3). ‘n.d.’ indicates not detected. B, Expression of *MEB1*, *MEB2*, and *MEB3* in response to jasmonic acid (JA) treatment. The chart shows RNA levels of rosette leaves and roots incubated with 50 µM JA for 1 day. Total RNA was subjected to qRT-PCR analysis. The transcript level of each gene was normalized relative to that of *UBQ10*. Data represent mean ± SE (*n* = 4).

Treatment with JA, a plant defense hormone, increases the number of ER bodies in Arabidopsis (Ogasawara et al. 2009; Geem et al. 2019; Stefanik et al. 2020). Therefore, we hypothesized that the ER-body-related *MEB1* and *MEB2* expression would be increased by JA, while the non-ER-body-related *MEB3* expression may not respond to the JA treatment. To test this hypothesis, we examined the expression of *MEB1*, *MEB2*, and *MEB3* in Arabidopsis roots and leaves with and without JA treatment (Figure 4B). In rosette leaves, JA treatment induced the expression of all three *MEB* genes; however, the enhancement of *MEB3* expression was much lower than that of *MEB1* or *MEB2*. In roots, all three *MEB* genes were expressed before the JA treatment; however, after the JA treatment, the expression of *MEB1* and *MEB2* was slightly enhanced, whereas that of *MEB3* was not affected. Thus, the gene regulatory mechanism of *MEB3* differs from that of *MEB1* and *MEB2*, suggesting that MEB3 is not strongly related to ER bodies induced by the JA treatment.

### *MEB3* expression is downregulated by both iron deficiency and supplementation

The *IRT1* gene, which encodes a plasma membrane-localized iron transporter, is upregulated by iron depletion and downregulated by iron supplementation (Vert et al. 2002; Connolly et al. 2002). Therefore, we examined changes in the expression levels of *MEB1*, *MEB2*, *MEB3*, *VIT1*, and *IRT1* both in the presence and absence of iron (Figure 5). Five-day-old wild-type (WT) seedlings grown in 100 µM iron were transferred to iron-deficient (0 µM) or -sufficient (200 µM) medium for 3 and 7 days, and total RNA was isolated from each sample to examine gene expression. Consistent with previous reports (Vert et al. 2002; Connolly et al. 2002), *IRT1* expression was strongly upregulated in iron-deficient medium and downregulated in iron-supplemented medium (Figure 5). By contrast, the expression of both *MEB3* and *VIT1* genes was downregulated in both iron-deficient and -sufficient media. The expression level of *MEB1* was similar under both iron-deficient and -sufficient conditions, and the expression of *MEB2* was upregulated under iron-sufficient and prolonged iron-deficient conditions. These results indicated that the expression regulation of vacuolar iron transporter genes (*MEB3* and *VIT1*) was different from that of the plasma membrane iron transporter gene (*IRT1*) and ER body iron transporter genes (*MEB1* and *MEB2*), suggesting that the mechanism of expression regulation of iron transporter genes is associated with the subcellular localization of the encoded proteins.

**Figure 5.**
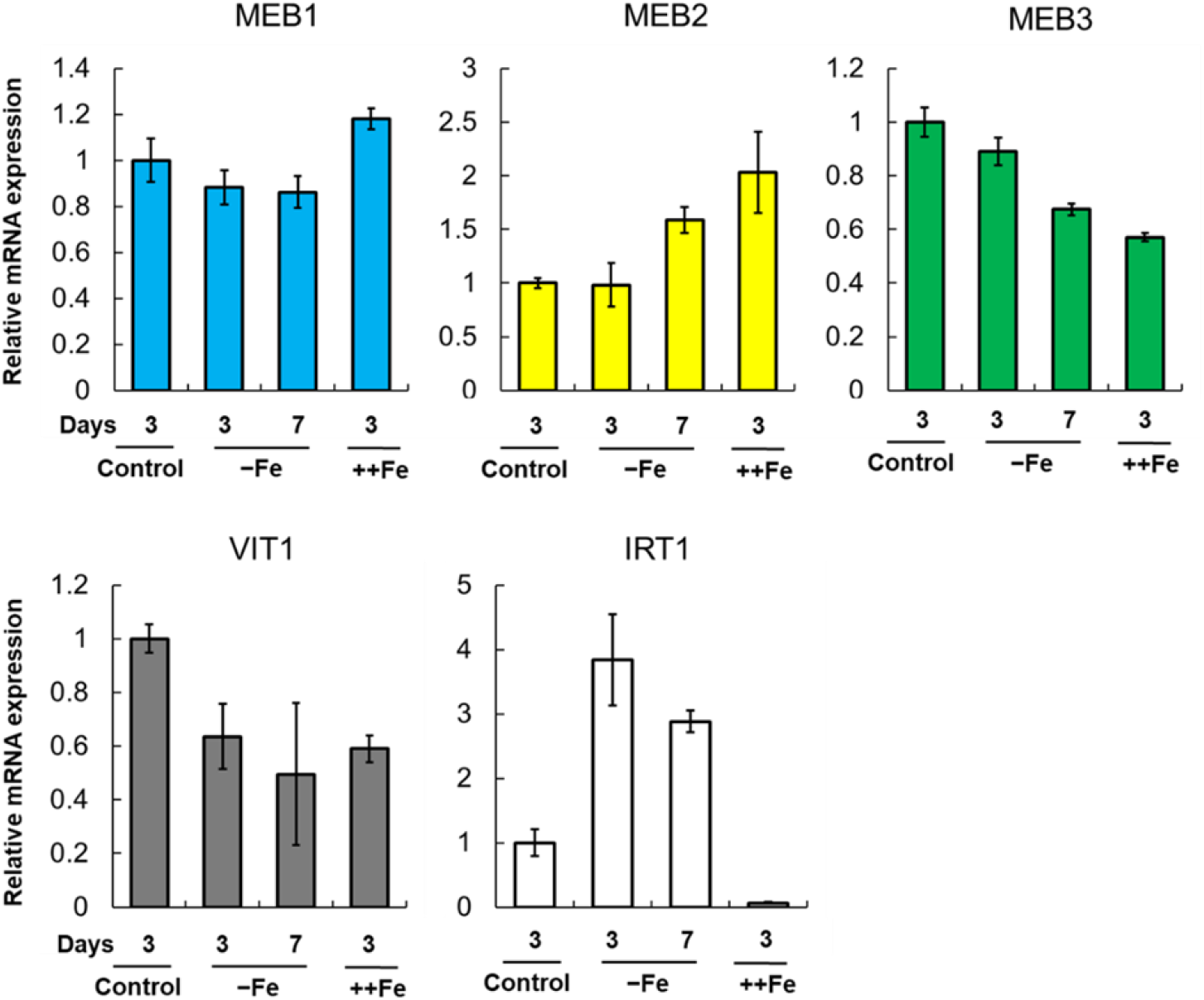
*MEB1*, *MEB2*, *MEB3*, *VIT1*, and *IRT1* gene expression under different iron concentrations. Plants were germinated in the normal medium (100 µM iron) for 5 days and then transferred to the normal (Control), iron-deficient (-Fe), or iron-sufficient (++Fe, 200 µM iron) medium for 3 or 7 days. The transcript levels of each gene are normalized to the *UBQ10* transcript levels. The error bars indicate SE (n = 4).

### *meb3* mutants exhibit reduced iron accumulation in roots

To examine the iron transport function of MEB3 in plants, we obtained two T-DNA insertion knockout mutants, *meb3-1* and *meb3-2*, in which the T-DNA was inserted in the first intron of *MEB3* (Figure 6A). Reverse transcription PCR (RT-PCR) analysis revealed no *MEB3* transcript in *meb3-1* and *meb3-2* mutants, indicating gene knockout (Figure 6B). Next, we measured iron contents of WT and *meb3* mutant plants grown on half-strength Murashige and Skoog (1/2 MS) medium containing 50 µM iron (normal condition). Consistent with the higher expression of *MEB3* in cotyledons and mature plant roots (Figure 4A), iron accumulation was higher in WT seedlings (Supplemental Figure 2) and roots of 21-day-old WT plants than in their *meb3* mutant counterparts (Figure 6C), suggesting that MEB3 is important for iron accumulation in plants. On the contrary, no reduction in manganese and zinc levels was observed in the roots and seedlings of *meb3* mutants compared with the WT counterparts (Supplemental Figures 2 and 3). Plants absorb iron from the soil, store it in roots, and distribute it to shoots via long-distance transport. To determine whether MEB3 is responsible for root-to-shoot iron distribution in response to iron availability, we transferred 5-day-old WT and *meb3* mutant seedlings from the germination medium (50 µM iron) to iron-sufficient (100 µM) or -deficient (0 µM) growth medium, and measured root and leaf iron levels after 21 days. In WT plants, root iron level was remarkably higher on iron-sufficient medium than on iron-deficient medium, whereas leaf iron level was only modestly higher on iron-sufficient medium than on iron-deficient medium (Figure 6D, Supplemental Figure 4). Leaf iron levels were not changed dramatically irrespective to the iron conditions and kept low levels compared to the root iron levels (Figure 6D). These results indicate that roots act as an iron storage organ, and root iron storage buffers leaf iron levels. We observed a similar trend of leaf iron-level-changes in *meb3* mutants, but surprisingly, the leaf iron level in *meb3* mutants was lower on iron-deficient medium but higher on iron-sufficient medium compared with the WT. Consistent with previous observations (Figure 6C), the iron level in *meb3* roots was lower than in WT root on iron-sufficient medium (Figure 6D). These findings indicate that the iron storage capacity, and consequently buffering function, of roots was reduced in the *meb3* mutants. This suggests that MEB3 contributes to iron translocation across organs in Arabidopsis.

**Figure 6.**
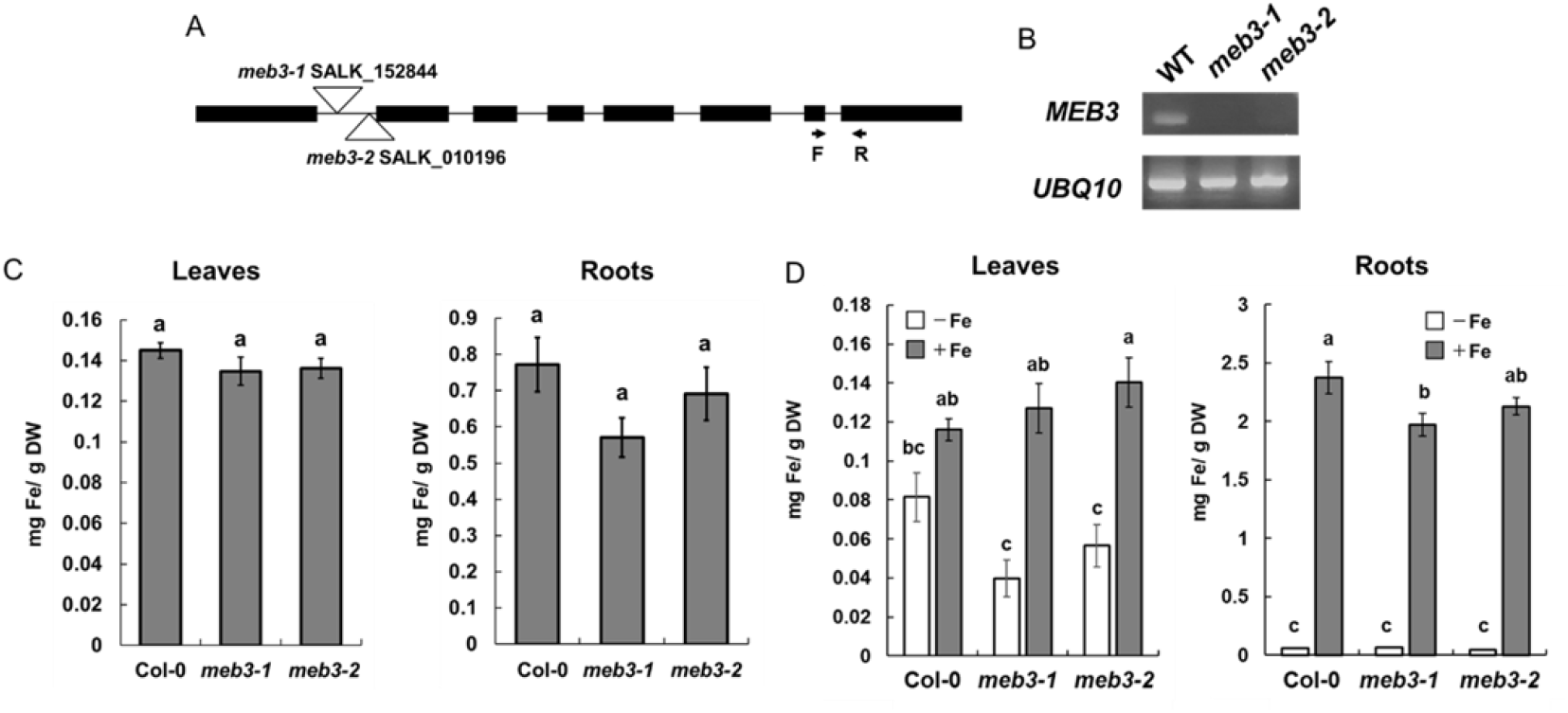
Loss of MEB3 affects the iron content in Arabidopsis leaves and roots. A, T-DNA insertion site in *meb3-1* (SALK_152844) and *meb3-2* (SALK_010196). The arrows indicate the binding sites and orientations of forward (F) and reverse (R) primers, which were used to determine *MEB3* transcript levels in the mutants by RT-PCR. B, Expression analysis of *MEB3* and *UBQ10* in the wild type (WT) and *meb3* mutants by RT-PCR. C and D, Quantification of leaf (left) and root (right) iron contents of 14-day-old wild-type, *meb3-1*, and *meb3-2* plants in 50 µM iron (C) or iron-sufficient (+Fe; 100 µM iron) and iron-deficient (-Fe; 0 µM) medium (D) by ICP-MS (C) or iron colorimetric assay (D). Five-day-old seedlings were transferred from medium containing 50 µM iron to iron-sufficient (+Fe; 100 µM iron) and iron-deficient (-Fe; 0 µM) medium and grown for 16 days before the iron contents were measured (D). Data represent mean ± SE (*n* = 10 in C, and 3 in D). Different letters above the columns indicate significant differences based on Tukey’s test (*p* < 0.05).

### *meb3* mutants exhibit reduced root and shoot growth under iron-deficient conditions

Because the iron storage capacity of *meb3* mutant roots was reduced, we hypothesized that the growth of *meb3* mutant plants is affected under iron-deficient conditions. To test this hypothesis, we transferred 5-day-old WT and *meb3* mutant seedlings from the germination medium (50 µM iron) to iron-sufficient (100 µM) or -deficient (0 µM) growth medium, and measured primary root length, shoot fresh weight, and total chlorophyll level after cultivation for 21 days. The primary root lengths of WT, *meb3-1*, and *meb3-2* plants were almost similar on iron-sufficient medium, but the primary root lengths of *meb3* mutants were reduced compared with those of WT plants on iron-deficient medium (Figure 7A, 7B). Similar trends were observed for shoot fresh weight (Figure 7A and 7C) and total chlorophyll content (Supplemental Figure 5) on iron-sufficient and -deficient medium. These results indicate that the knockout mutation of *MEB3* mildly reduces plant growth under iron-deficient conditions, suggesting that MEB3 contributes to iron storage to enable shoot and root growth.

**Figure 7.**
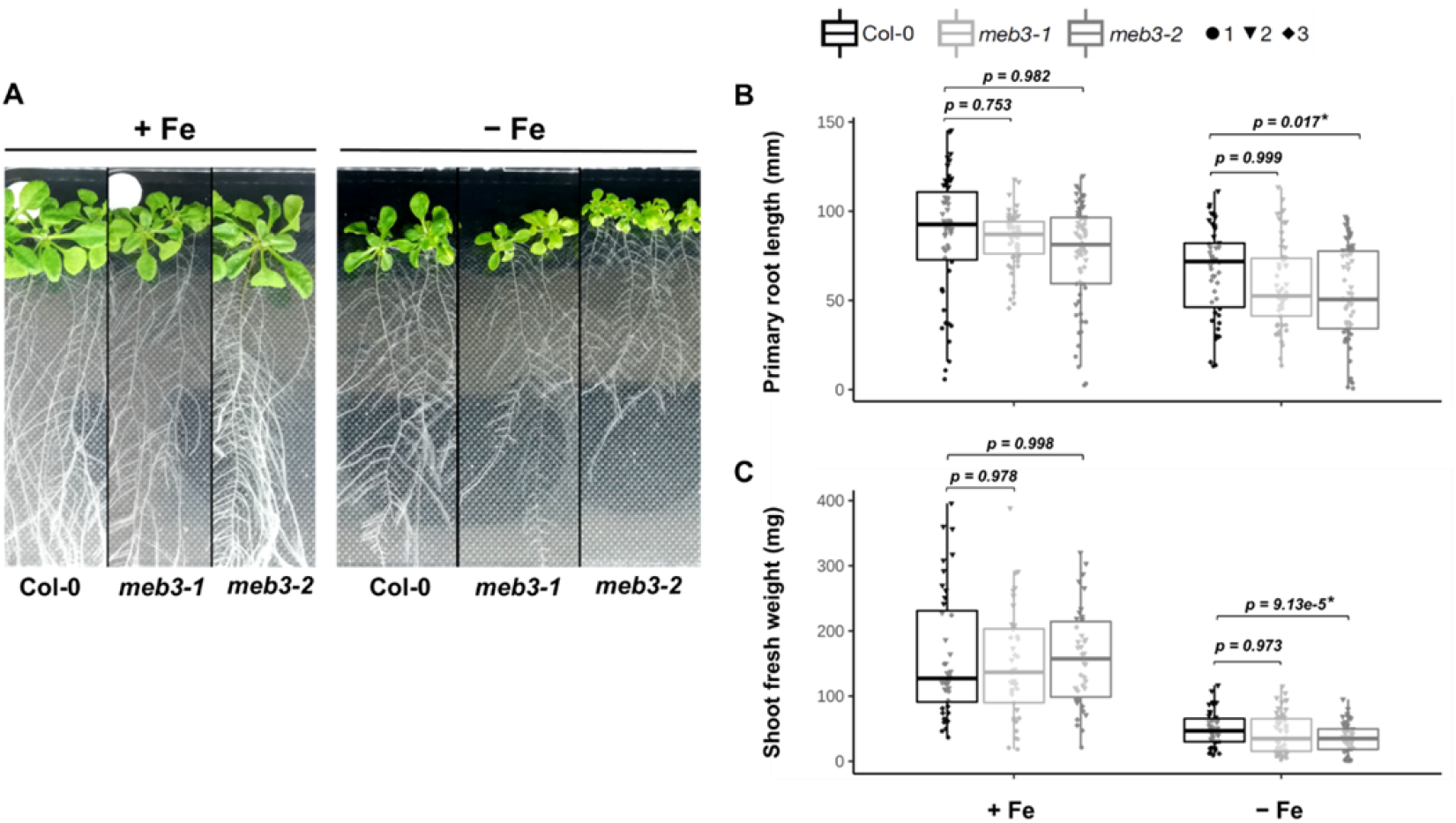
Growth phenotype of *meb3* knockout mutants. A, Photograph of 21-day-old wild-type (WT), *meb3-1*, and *meb3-2* plants. Five-day-old seedlings (grown in ½ MS including 50µM Fe-EDTA) were transferred to 100 µM Fe-EDTA or Fe-free medium and grown for 16 days. B and C, Primary root length (B) and shoot fresh weight (C) of WT, *meb3-1*, and *meb3-2* plants treated as the same as in (A). Asterisks indicate significant differences between WT and mutant plants. Circle, triangle, and diamond represent values of biological replicates. The *p*-values displayed on the boxplot were obtained using Tukey’s test applied over log2-transformed data.

## DISCUSSION

### Arabidopsis MEB3 is a vacuolar iron transporter

Arabidopsis MEB3, like MEB1 and MEB2, showed iron transport activity when expressed in yeast. However, we found that MEB3 localized to the tonoplast, but not to ER bodies, in Arabidopsis leaves and roots. Thus, our results suggest that unlike MEB1 and MEB2, which function as ER body-localized iron transporters (Yamada et al. 2013), Arabidopsis MEB3 functions as a vacuolar iron transporter.

In budding yeast, the vacuolar iron transporter CCC1 controls cytosolic iron level, and the *ccc1* mutant does not grow in iron-containing medium, because of cytosolic iron toxicity (Kim et al. 2006). Overexpression of Arabidopsis *VIT1* and *MEB1–3* complemented the growth inhibition phenotype of the *ccc1* mutant (Figure 2B and 2C), indicating that VIT1 and MEB1–3 proteins act as iron transporters and remove iron from the cytosol. However, the growth of *MEB1*- and *MEB2*-expressing *ccc1* mutant cells was lower than that of *VIT1*- and *MEB3*-expressing yeast, suggesting that the iron transport function of MEB1 and MEB2 is less strong than that of VIT1 and MEB3. Additionally, we found that MEB1 and MEB2 localized to the ER, while VIT1 and MEB3 localized to the tonoplast in yeast cells (Figure 2A). Therefore, the functional difference between MEB1/MEB2 and VIT1/MEB3 could be attributed to the difference in their subcellular localization patterns rather than the difference in their transporter activities; the iron accumulation capacity of vacuoles is higher than that of the ER.

Amino acid residues crucial for the iron transport activity of EgVIT1 (Kato et al. 2019) were conserved in the VIT1 subfamily, but some of these residues were not conserved in the SEN1/NODULIN21 and MEB subfamilies (Supplementary Figure 1). Nevertheless, Gollhofer et al. (2014) showed that two SEN1/NODULIN21 subfamily proteins, namely, Arabidopsis VTL1 and VTL2, exhibit iron transport activity. Consistently, our data indicated that Arabidopsis MEB1, MEB2, and MEB3 are involved in iron transport (Figure 2B). The SEN1/NODULIN21 subfamily proteins regulate iron transport into the symbiosome in the root nodules of soybean and alfalfa plants (Brear et al. 2020; Liu et al. 2020; Walton et al. 2020). These findings indicate that the conservation of these amino acid residues is not crucial for the iron transport activity of SEN1/NODULIN21 and MEB subfamily proteins, which suggests that their iron transport mechanism is unique.

Because our results suggest that Arabidopsis MEB3 is a vacuolar protein, implying the existence of a mechanism that distinguishes between the subcellular localization of MEB3 and MEB1/2 in Arabidopsis. Surprisingly, the *B. napus* ortholog of MEB2 localizes to the tonoplast but not to ER bodies (Zhu et al. 2016), indicating that the subcellular localization of MEB proteins is not conserved even within the Brassicaceae family. The vacuolar transport signal of MEB3 and BnMEB2 remains to be identified. Wang et al. (2014) showed that VIT1 possess a dileucine motif (D/EXXXLL), which leads to the vacuolar targeting of VIT1. This motif also exists in NRAMP3 and NRAMP4 and is responsible for their vacuolar localization (Müdsam et al. 2018). However, we could not find the D/EXXXLL motif in MEB3. The MEB subfamily proteins harbor a long motif in the N-terminal region, which is more varied in sequence than the transmembrane domain in their C-terminal region (Yamada et al. 2013). This uncharacterized motif in the N-terminal region might determine the subcellular localization of MEB subfamily proteins in plants.

### MEB3 regulates iron accumulation in roots

The expression level of *MEB3* was higher in cotyledons and roots than in other tissues (Figure 4A), indicating that MEB3 functions in cotyledons and roots. Additionally, the root iron content of *meb3* mutants was lower than that of the WT, indicating that the absence of MEB3 was responsible for the reduced vacuolar iron level roots in *meb3* mutants (Figure 6C). These findings suggest that MEB3 regulates iron accumulation in Arabidopsis roots. Treatment with JA, a defense-related hormone known to induce ER body formation (Stefanik et al. 2020), modestly induce *MEB3* expression in leaves (Figure 4B). By contrast, the expression of *MEB1* and *MEB2*, which encode ER body-localized proteins (Yamada et al. 2013), was strongly induced by JA treatment (Figure 4B). This suggests that the function of MEB3 *in planta* is different from that of MEB1 and MEB2; for example, MEB3 may not be involved in plant defense, unlike MEB1 and MEB2.

Besides Arabidopsis MEB3, the SEN1/NODULIN21 subfamily proteins are also involved in root vacuolar iron accumulation (Rodríguez-Haas et al. 2013; Ram et al. 2021). Arabidopsis *vlt3* and *vtl5*/*nodulin-like21* mutants exhibit reduced root iron contents compared to the WT (Gollhofer et al. 2011). Arabidopsis FPN2 is a vacuolar iron transporter that does not belong to the VIT family but is expressed in roots (Morrissey et al. 2009). Therefore, one can expect that these proteins and MEB3 regulate vacuolar iron contents under iron-sufficient conditions in a functionally redundant manner, which may be why the *meb3* mutants do not show a complete loss of root iron.

The flexibility of root iron accumulation helps plants to grow under iron stress conditions. Expression of the plasma membrane iron transporter gene *IRT1* was suppressed in Arabidopsis seedlings and roots in medium containing excess iron (Figure 5), suggesting that plants try to reduce unwanted iron uptake from the medium. A similar phenomenon was observed in rice (*Oryza sativa*) in previous studies, where *OsIRT1* expression was suppressed in roots under excess iron conditions (Aung et al. 2018; Aung and Masuda 2020). By contrast, the expression of two vacuolar iron transporter genes, *OsVIT2* and *BnMEB2*, was upregulated in roots under excess iron conditions (Aung et al. 2018; Aung and Masuda 2020; Zhu et al. 2016), suggesting that plants increase the abundance of vacuolar iron transporters to enhance the sequestration of cytosolic iron in vacuoles. In the current study, Arabidopsis *MEB3* and *VIT1* genes did not respond to excess iron (Figure 5), but we could not exclude the possibility that MEB3 and VIT1 do not participate in the vacuolar sequestration of cytosolic iron under excess iron conditions. The iron content of leaves seemed to be slightly increased in *meb3* mutant plants grown on iron-sufficient medium (Figure 6D). Because of the low iron accumulation capacity of *meb3* roots, excess cytosolic iron in roots might be translocated to shoots in *meb3* mutants.

### MEB3 is involved in the root-to-shoot translocation of iron

We showed that MEB3 is involved in root iron accumulation, and that the root iron level of *meb3* mutants was lower than that of the WT in normal medium. Although *MEB3* expression was not high in shoots, the shoot iron levels decreased and plant growth reduced in *meb3* mutants compared with WT plants upon transfer from the normal medium to iron-deficient medium (Figure 7C). These findings suggest that root iron accumulation promotes shoot growth, presumably by the mobilization of root iron reserves to the shoot under iron-deficient conditions. In Arabidopsis, it has been postulated that vacuolar NRAMP3 and NRAMP4 proteins are involved in unloading iron from the vacuoles into the cytosol (Lanquar et al. 2005), and the unloaded cytosolic iron chelates with NA (Klatte et al. 2009). The resultant Fe^2+^-NA complex is captured by secretory vesicles via nitrate transporter 1/peptide transporter family proteins NRT5.8 and NRT5.9 and then secreted into the phloem for long-distance transport to the shoot (Chen et al. 2021; Chao et al. 2021). The expression of genes involved in the iron remobilization pathway, such as *NRAMP3*, *NRAMP4*, NA biosynthesis, *NRT5.8*, and *NRT5.9*, is enhanced in response to iron deficiency (Lanquar et al. 2005; Klatte et al. 2009). By contrast, the expression of *MEB3* and *VIT1* was reduced (Figure 5). This finding is consistent with the results of Gollhofer et al. (2011), who showed that the expression of *VTL1* and its homologs is reduced in roots in response to iron deficiency. Therefore, it is plausible that vacuolar iron influx is reduced, and vacuolar iron remobilization is increased, in roots for efficient iron translocation to the shoot under iron-deficient conditions.

In iron-sufficient medium, root iron accumulation was reduced, but shoot iron levels were enhanced in the *meb3* mutant, indicating that the root-to-shoot transport of iron increases in *meb3* mutants under iron-sufficient conditions. This finding suggests that the MEB3-mediated iron storage capacity in the root is responsible for buffering leaf iron levels against environmental changes of iron levels. Therefore, MEB3-mediated vacuolar iron storage in Arabidopsis roots has two functions: 1) storage of iron to prepare for iron shortage in the shoot and 2) buffering shoot iron levels by storing iron in the root.

Proper iron distribution is essential for plant growth. For example, Arabidopsis *vit1* mutant shows no difference in iron contents relative to the WT but exhibits impaired iron accumulation in provascular strands in seeds (Kim et al, 2006). The unusual distribution of iron in the *vit1* mutant affects seedling development under iron-deficient conditions (Chu et al. 2017). Similarly, we found that MEB3 is involved in the proper distribution of iron between shoots and roots, and its absence affects plant growth. Considering that MEB subfamily proteins exist in almost all plant species in addition to the ER body-producing plants, it is suggested these proteins are involved in similar functions (i.e., vacuolar iron accumulation) to promote plant growth.

## MATERIALS AND METHODS

### Plant materials, plant growth condition, and yeast strains

*Arabidopsis thaliana* ecotype Columbia (Col-0) was used as the WT in this study. Arabidopsis T-DNA insertion mutants *meb3-1* (SALK_152844) and *meb3-2* (SALK_010196) were obtained from the Arabidopsis Biological Resource Center (ABRC; OH, USA). Seeds were surface-sterilized with 70% (v/v) ethanol and germinated photoautotrophically at 22°C under continuous light (approximately 100 µE s^-1^ m^-2^) on 1/2 MS medium (half-strength MS basal salt mixture [2623020; MP Biomedicals], 1% [w/v] sucrose, and 0.5% [w/v] MES-KOH [pH5.7]) containing 0.4% (w/v) Gellan Gum (Wako). For iron-deficient experiments, after 5 days, the seedlings were transferred to iron-sufficient medium (1/2 MS medium supplemented with100 µM NaFe(III)-EDTA) or iron-deficient medium (1/2 MS medium supplemented 100 µM Na_2_-EDTA instead of NaFe(III)-EDTA). Then, eight seedlings from each treatment were placed in separate square plates and incubated in a growth chamber for 16 days at 22°C under continuous light. Primary root length was measured using the ImageJ software. *Nicotiana tabacum* Bright Yellow 2 (BY-2) cell suspension cultures were grown in MS basal salt mixture (M5524; Sigma) supplemented with 3% (w/v) sucrose, 0.2% (w/v) KH_2_PO_4_, 0.2 mg/L 2,4-dichlorophenoxyacetic acid, 1 mg/L thiamine, and 100 mg/L myo-inositol (pH 5.5) in the dark at 25°C on an orbital shaker (180 rpm). The *CCC1*-deficient yeast strain (Δ*ccc1*::KanMX4; BY4741 background) was used for the heterologous expression of Arabidopsis *MEB1*, *MEB2*, *MEB3*, and *VIT1* cDNAs, to examine the subcellular localization patterns and iron transport activities of the encoded proteins.

### Plasmid construction and plant transformation

To examine the subcellular localization of MEB3, a cDNA fragment of *MEB3* with and without stop codon was amplified and cloned into the Gateway Entry vector pENTR/SD/D-TOPO (Thermo Fisher Scientific). The inserted gene was transferred into the destination vectors pGWB405 and pB5tdGW (gifts from T. Nakagawa), harboring *GFP* and *tdTOM* reporter genes, respectively (Nakagawa et al., 2007), using Gateway LR Clonase Enzyme mix (Thermo Fisher). The resultant constructs pB5tdGW/tdTOM-MEB3 and pGWB405/GFP-MEB3, carried *Prom35S*:*GFP-MEB3* and *Prom35S*:*tdTOM-MEB3*, respectively. The pB5tdGW/tdTOM-MEB3 construct were transformed into Col-0 plants via *Agrobacterium*-mediated transformation using the floral dip method (Clough and Bent 1998). To perform iron transport assay in yeast cells, full-length *VIT1* cDNA was amplified from the Arabidopsis cDNA library and subcloned into pENTR1A (Thermo Fisher) with *Sal*I and *Eco*RV. The DNA fragments containing *MEB3* (pENTR-MEB3) and *VIT1* (pENTR-VIT1) were cloned into the p415 GAL1-GW vector (Yamada et al. 2013) via the LR reaction to generate p415/MEB3 and p415/VIT1 vectors, respectively. To examine the subcellular localization of MEB1–3 proteins in yeast cells, full-length *MEB1*, *MEB2*, and *MEB3* cDNA were cloned into pAG416GAL-EGFP-ccdB (Addgene plasmid # 14315) vector via the LR reaction.

### Transient expression assays using Arabidopsis protoplasts

Protoplasts were isolated from 20 rosette leaves of 3-week-old plants using the tape-sandwich method described by Wu et al. (2009). The peeled leaves were gently shaken with enzyme solution for 90 min at room temperature. The pGWB405/GFP-MEB3 was transiently expressed in protoplasts using polyethylene glycol (PEG)-mediated transfection (Yoo et al., 2007). The transfected protoplasts were incubated in the plant growth room (22°C, 16 h light/8 h dark cycle) for 1 day, harvested by centrifugation at 100 × *g*, and subsequently observed under a microscope. To isolate protoplasts from BY-2 cells, 1.5 g of 4-day-old cells was incubated in 10 mL enzyme solution (1% [w/v] cellulase Onozuka RS, 0.1% [w/v] Pectolyase Y-23, 0.4 M mannitol, 10 mM CaCl_2_, 5 mM MES-Tris [pH 5.5]) at 28°C for 1 h with gentle shaking.

### Confocal microscopy

A confocal laser scanning microscope (LSM880; Carl Zeiss) was used to observe fluorescent proteins. GFP was detected using an argon laser (488 nm) and a 505/530 nm band-pass filter, chlorophyll *b* was observed using a helium-neon laser (633 nm) and 640/720 nm band-pass filter, and tdTOM signal was observed using a 561 nm diode-pumped solid-state laser and 560/640 nm band-pass filter.

### Measurement of chlorophyll content

The total chlorophyll content of 21-day-old plant shoots was measured according to the method described by Harbort et al. (2020). The absorbance of 100 µL samples was measured at 652 nm using a microplate reader (Infinite M200 Pro, Tecan).

### Iron quantification

The total iron content of Arabidopsis leaves and roots was quantified either using the iron colorimetric assay (Gao et al. 2021) or by inductively coupled plasma-mass spectrometry (ICP-MS) (Almario et al. 2017). To measure iron content by the colorimetric assay, leaf and root samples were dried at 80°C for 20 h, and their dry weights were measured. To measure iron content by ICP-MS, 7-day-old seedlings and the leaves and roots of 21-day-old plants grown on 1/2 MS medium (50 µM Fe) were freeze-dried at -20°C overnight. The dried plant material was weighed and homogenized to a fine powder. Approximately 5 mg of the powdered plant material was digested with 500 μL of 67% (w/w) HNO_3_ in 15 mL Falcon tubes overnight at room temperature. Then, loosely closed tubes containing the samples were placed in a 95°C water bath until the liquid was completely clear (approximately 30 min). After being cooled to room temperature for 10–15 min, the samples were placed on ice, and 4.5 mL of deionized water was carefully added to the tubes. The samples were then centrifuged at 2,000 × *g* for 30 min at 4°C, and the supernatants were transferred into new tubes. The elemental concentration of iron was determined using Agilent 7700 ICP-MS (Agilent Technologies).

### Iron transport assay

The iron resistance assay was performed in yeast cells as described previously (Yamada et al. 2013). Briefly, 5 µL of yeast cell suspension (OD_600_ = 0.1 or 0.01) was spotted on SGal-agar medium containing 3 mM ammonium Fe(II) sulfate.

To perform iron uptake assay, yeast strains were grown overnight in YPD medium and then washed once with water. The washed cells were suspended in water and applied to 100 mL of SGal medium containing 0.5 mM ammonium Fe(II) sulfate. Cells were grown for 16 h at 30°C and then centrifuged at 5,000 rpm for 5 min. The precipitated cells were washed twice with 1 mM EDTA and once with water, and suspended in Milli-Q water. Subsequently, 1 mL of cells (OD_600_ = 10) was transferred into 1.5 mL tubes. After centrifugation at 10,000 rpm for 5 min, the cells are dried completely at 60°C for 3 days. The total iron content of yeast cells was measured by the iron colorimetric assay.

### JA treatment

The leaves and roots of 2-week-old WT plants were cut and placed on the surface of water containing 50 μM methyl jasmonate (MeJA; Sigma-Aldrich, St. Louis, USA). The JA treatment was performed for approximately 24 h at 22°C under continuous light.

### RNA isolation and quantitative real-time PCR (qRT-PCR)

To analyze mRNA expression levels in each plant organ, total RNA was isolated from 5-day-old cotyledons and the rosette leaves and roots of 14-day-old plants. After 2 weeks of germination, plants were transferred to soil and grown for 4 weeks. Then, total RNA was isolated from the cauline leaves, stems, green siliques, and flowers of 6-week-old plants. Isolation of total RNA from leaves and roots was performed using the TRIzol Reagent (Molecular Research Center, Cincinnati, USA), and isolation of total RNA from stems, green siliques, and flowers was performed according to the method described by Oñate-Sánchez and Vicente-Carbajosa (2009). The isolated total RNA was resuspended in distilled water and treated with DNase I (Thermo Fisher). Then, first-strand cDNA was synthesized from 2 µg of total RNA using Ready-to-Go RT-PCR beads (GE healthcare) and an oligo(dT) primer. The expression of *MEB1*, *MEB2*, *MEB3*, and *VIT1* was analyzed by qRT-PCR (QuantStudio 6, Thermo Fisher) using TaqMan Real-Time PCR Assays (Thermo Fisher), according to the manufacturer’s instructions. The transcript levels of *MEB1*, *MEB2*, *MEB3*, *VIT1*, and *IRT1* genes under iron-deficient conditions were analyzed using PowerUp SYBR Green Master Mix (Thermo Fisher). The gene-specific primer sets were designed with Primer3Plus software, and *UBQ10* was chosen as the housekeeping gene. The expression of target genes was normalized relative to the constitutive expression of *UBQ10* using the amplification efficiency calibrated calculation method.

## ACKNOWLEDGMENTS

We thank Tsuyoshi Nakagawa (Shimane University) for sharing vectors, Justyna Łabuz and Paweł Hermanowicz (Jagiellonian University) for their assistance with the confocal microscope, Sabine Ambrosius (University of Cologne) for technical assistance, and the Biocenter MS Platform Cologne for the ICP-MS measurements.

## AUTHOR CONTRIBUTIONS

K.E. and K.Y. designed the research and wrote the manuscript. K.E and A.K.B., and M.M. performed the experiments. K.E and A.K.B designed experiments and analyzed the data. K. E. and A.W. generated transgenic yeast and plant lines. S.K. performed the ICP-MS analyses.

## FIGURE LEGENDS

Supplementary Figure 1. Aliment of DUF125 domains, including key amino acid residues required for iron transport activity. Black open boxes indicate amino acid residues important for the iron transport activity of EgVIT1 (i.e., D43, E72, M80, and Y175) (Kato et al. 2019). The orange-colored open box indicates the Gly76 residue of EgVIT1, which is crucial for iron transport activity (Mary et al. 2015). The proteins used in the analysis are listed in Supplementary Table S1. ‘-’ indicates deleted amino acids.

Supplementary Figure 2. Iron, manganese, and zinc contents of 7-day-old seedlings grown on the 1/2 MS medium containing 50 µM Fe. The metal contents of dried plants were measured by ICP-MS. Data represent mean ± SE (*n* = 12 Col-0 seedlings and 10 *meb3-2* seedlings). Asterisks above the columns indicate significant differences based on Student’s *t*-test (p < 0.05).

Supplementary Figure 3. Manganese and zinc contents of the leaves and roots of 14-day-old plants grown on 1/2 MS medium containing 50 µM Fe. Plants used in this experiment are the same as those shown in Figure 6C experiment. Data represent mean ± SE (*n* = 10).

Supplementary Figure 4. Magnified chart of root iron contents of plants grown on iron-deficient medium, as shown in Figure 6D. Data represent mean ± SE (*n* = 3).

Supplementary Figure 5. Chlorophyll contents of wild-type (WT), *meb3-1*, and *meb3-2* plants treated as the same as in Figure 7. Asterisks indicate significant differences between WT and mutant plants. Circle and square represent values of biological replicates (*n* = 10). The *p*-values displayed on the boxplot were obtained by Tukey’s test.

Supplementary Table S1. List of proteins used to construct the phylogenetic tree shown in Figure 1

Supplementary Table S2. Summary statistics for the mesurements shoot fresh weight and primary root length

Supplementary Table S3. List of primers used in this study

